# Optimization of accelerated solvent extraction of fatty acids from *Coix* seeds using chemometrics methods

**DOI:** 10.1101/496935

**Authors:** Xing Liu, Kai Fan, Wei-Guo Song, Zheng-Wu Wang

## Abstract

This study investigated the optimization of accelerated solvent extraction (ASE) of fatty acids (FAs) from three *Coix* seeds (small *Coix* seed, SCS; big *Coix* seed, BCS; translucent *Coix* seed, TCS) by chemometrics methods. Partial least-squares regression (PLSR) and backpropagation neural network (BPNN) were applied to build models that reflect the relationship between content of FAs and extraction conditions (temperature, time, and extraction solvent). Genetic algorithms (GAs) and particle swarm optimization (PSO) were utilized to optimize the combination of extraction conditions. The composition of FAs was analysed by gas chromatography-mass spectrometry (GC-MS). The PLSR models could reflect the relationship of FA content in both BCS and SCS and extraction conditions well, while the BPNN model was more suitable for TCS. The optimal extraction conditions for BCS and SCS were obtained by GAs, whereas those of TCS were obtained by PSO. The FA compositions of the three *Coix* seeds exhibited differences. The results show that ASE combined with chemometrics methods can rapidly and effectively obtain the optimal conditions for the extraction of FAs from *Coix* seed and there are differences in the extraction conditions and compositions of FAs among different varieties of *Coix* seed, but all the extraction time is shorter than other extractions methods.

## Introduction

*Coix* seed is the mature kernel of *Coix lachryma-jobi* L., a grain crop in the Gramineae family, and has long been used as a traditional Chinese medicinal herb and food source. *Coix lachryma-jobi* L. is widely distributed in China, Thailand, Burma, Korea, Japan, and Brazil [1]. There are many reported pharmacological and physiological effects of *Coix* seed, including anti-tumour [2], anti-inflammatory [3], anti-allergic [4], and immunoregulation [5]. These effects result from diverse biologically active components in *Coix* seed [6, 7], which mainly exist in *Coix* seed oil [8], such as coixenolide, coixol, and sterols. *Coix* seed oil is mainly composed of the fatty acids (FAs), and the content of FAs can reflect the yield of the extracted oil and the content of other active ingredients to a certain degree. Moreover, the kinds of FAs have an important impact on the nutritive value of *Coix* seed oil. Due to its many benefits, it is reasonable to pursue the optimization of the extraction conditions of *Coix* seed oil. The common extraction techniques, such as Soxhlet extraction [9], microwave extraction [10], sonication extraction [11], and supercritical fluid extraction [8], are time-consuming and/or complex. The extraction yield of FAs from *Coix* seed is dependent on the following factors: temperature, time, pressure, extraction solvent, particle size, and solid-liquid ratio. Accelerated solvent extraction (ASE), an extraction procedure using organic solvents at high temperatures (elevated temperatures up to 200 °C) and pressures (up to 3000 psi) above the boiling point for shorter time (low to several minutes), can increase target compound solubility, solvent diffusion rate, and mass transfer. It can also decrease solvent viscosity and surface tension, which has been shown to be equivalent to the standard EPA extraction methodology (Method 3545) in terms of precision and recovery [12]. Furthermore, the extraction process of ASE has the advantage that needs less solvent, is automated and quick, and can retain the sample in an oxygen- and light-free environment [13]. Currently, many studies have applied the ASE method to extract lipids and FAs from cereal, egg yolk, fish, fish tissue, and chicken muscle [14-17]. It has also been reported that the FA composition was not affected by the extraction temperature of ASE [18].

Our previous study has proven that the extraction yield of crude fat in *Coix* seed by ASE is not lower than that of Soxhlet extraction or sonication-assisted supercritical fluid extraction [11, 19]. However, the extraction process remains to be optimized from the perspective of energy-saving, and there is limited information on how to optimize the extraction conditions (temperature, time, and extraction solvent) of FAs from *Coix* seed by ASE. Thus, a full factorial design (FFD) was applied to design the experiment, chemometrics methods, partial least-squares regression (PLSR) and a backpropagation neural network (BPNN), were used to build the relationship between FAs and extraction conditions, and genetic algorithms (GA) and particle swarm optimization (PSO) were utilized to optimize the extraction conditions. The PLSR, as a chemometrics method and linear regression tool, is one of the most widely applied multivariate statistical data analysis method [20]. The BPNN is a classical domain-dependent technique for nonlinear system modelling. It is composed of an input layer, hidden layer, and output layer, and works by measuring the output error, calculating the gradient of this error, and adjusting the neural network weights (and biases) in the descending gradient direction. That is, the BPNN is a gradient-descent local search procedure that is expected to stagnate in local optima in complex landscapes [21]. The GA and PSO are most popular optimization algorithms, and they employ a population of individuals to solve the problem on hand [22]. It has been reported that GA, which are parallel randomly search optimization algorithms, can be successfully applied to identify global optimizations of multidimensional functions by selecting, crossover, and mutation operations [23]. The PSO is a stochastic evolutionary computation technique, inspired by the social behaviour of bird flocking [24]. Similar to GA, the PSO system is initialized with a population of random solutions and can search for optimum conditions by the updating of generations [25]. The content of oils is significantly difference due to the region in which the crop is grown as well as varietal diversity [19]. Therefore, it is necessary to study the FA content of different varieties of *Coix* seed. In the present study, temperature, time, and extraction solvent were optimized to extract FAs from different *Coix* seeds by ASE combined with chemometrics methods. The composition of FAs was determined by gas chromatography-mass spectrometry (GC-MS).

## Materials and methods

### Materials

Small *Coix* seeds (SCS, aspect ratio=0.23 cm: 0.26 cm) and big *Coix* seeds (BCS, aspect ratio=0.32 cm: 0.40 cm) were purchased in Anshun Municipality and Guizhou province, China, respectively; translucent *Coix* seeds (TCS, aspect ratio=0.21 cm: 0.22 cm) were purchased in Putian Municipality, Fujian province, China. The three categories of *Coix* seed were named in terms of their appearance and size and are shown in Fig.1. Before the experiment, defective granules were removed from all samples, and the seeds were ground until they could pass through a 425-μm mesh sieve. The sieved powders were used in subsequent analysis. All chemicals were purchased from the China National Pharmaceutical Group (Sinopharm, Beijing, China) and were of analytical grade.

**Fig 1.**
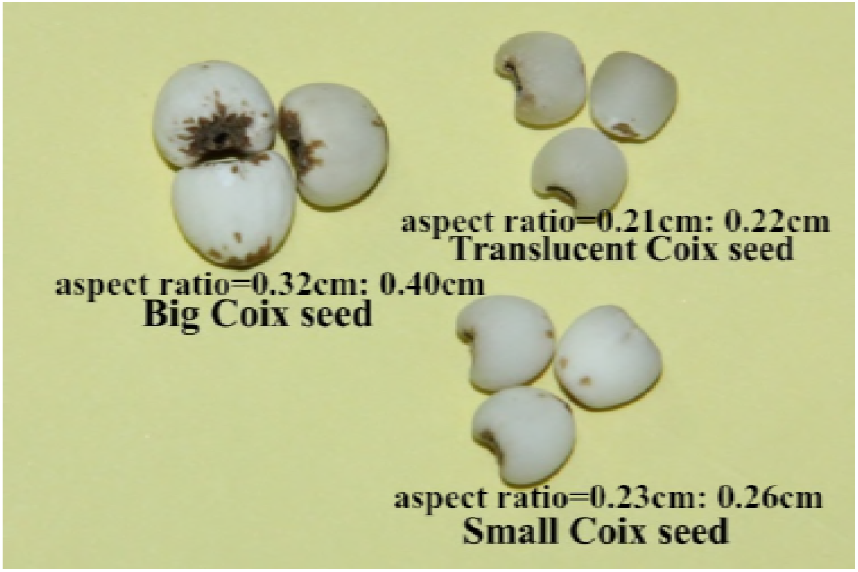
The appearance and size of the three categories of *Coix* seed.

### Crude fat extraction and FA determination

Crude fats and FAs of the three categories of *Coix* seed were extracted by an ASE apparatus (ASE 350, Thermo Fisher Scientific, Waltham, MA, USA). *Coix* seed samples of 10.0 g were weighed and poured into a 66-mL zirconium extraction cell with a cellulose filter in the cell outlet. The extraction cell was arranged in the cell tray, and the sample was extracted using a combination of conditions obtained from the FFD experimental design. The automated extraction cycle was as follows: the cell containing sample was prefilled with the degassed extraction solvent (acetone or petroleum ether), pressurized (1600 psi), and then heated. The cycle time varied with the change in temperature (100, 110, 120, or 130 °C). When the temperature was higher than the 130 °C, the colour of *Coix* seed oil became dark. The last step in the cycle was a static period (5, 10, 15, or 20 min). Then, the cell was rinsed with fresh extraction solvent (60% of the extraction cell volume) and purged with a stream of nitrogen. The extraction cycles were repeated twice. The oil was collected into glass vials and concentrated immediately by rotary evaporators (35 °C). The FA content in the concentrated solution was determined by the AOAC method (939.05) and expressed as mg of KOH required neutralizing FAs in 100 g *Coix* seed. The concentrated solution obtained in the optimal extraction condition was evaporated to dryness by water bath, and dried for 1 h at 100 °C ± 5 °C, then cooled down for 0.5h in the desiccator and weighed. The process above was repeated until achieved constant weight. This method was calculated the crude fat content.

### GC-MS analysis of FAs

One gram of *Coix* seed oil was dissolved in 40 mL of n-hexane, then 40 mL 0.4 M KOH-MeOH solution was poured into a test tube, which was vigorously shaken, and the mixture was placed for 30 min. After being fully saponified, 10 mL distilled water was put into the test tube, and the supernatant was used to analyse the composition of FAs by GC-MS (7890A-5975C Agilent, Santa Clara, CA, USA). The GC column was a DB-5ms (30 m × 0.25 mm × 0.25 µm). The carrier gas was helium (99.999%), and the flow rate was 1 mL/min. Both the injector temperature and detector temperature were 280 °C The program sequence of the column temperature was as follows: initial temperature 60 °C, held for 3 min, increased to 300 °C at 5 °C/min, and held for 14 min. The MS ion source was electron impact mode at an ionization voltage of 70 eV with an ion source temperature of 230 °C. The full-scanning range of MS was 33–500 amu. The results were obtained from the NIST 2011 mass spectral data base.

### Chemometrics methods and statistical analysis

The ASE experiment was designed to consider the factors of temperature (100, 110, 120, or 130 °C), extraction time (5, 10, 15, or 20 min) and extraction solvent (acetone or petroleum ether) and was carried out according to the FFD, whose total trial number was 32.

The Kennard-Stone algorithm was used to partition the calibration (75%) and validation sets (25%) [26], and the criterion was to select the samples one by one which was the furthest distance from each other in the group, namely, according to the Euclidean distance, so they could spread throughout the multivariate space. Linear models of the FAs extraction were established by PLSR. The latent variables of PLSR were determined by 10-fold cross-validation with the lowest root mean square error of cross validation (RMSECV). The performances of calibration set models were valued by the RMSECV and the coefficient of determination (R^2^), and validation set models evaluated by root mean square error of prediction (RMSEP) and R^2^ between predicted value and actual value. Nonlinear models of the FA extraction were built by a BPNN, whose input layer nodes were 3, hidden layer nodes were 10, and output layer node was 1. The BPNN models were estimated by RMSEP and the R^2^ of validation sets. In this study, the PLSR and BPNN models with the highest R^2^, as well as the lowest RMSECV and RMSEP, were considered as the optimal result.

Two extreme value searching algorithms, GAs and PSO, were applied to screen the optimum extraction conditions (extraction solvent, time, and temperature). Generally, a given problem can be regarded as an individual coded by chromosome strings in the GAs. The individual fitness function values, evaluating a chromosome about the objective function of the optimization problem, are used as the evaluation index of individual quality. In the process of population evolution, selection, crossover, and mutation are continuously applied to gradually reach optimal solutions until it generates the global optimal solution [27-29]. The parameters of the GAs were as follows: evolutional generation 100, population size 100, crossover probability 0.8, and mutation probability 0.6. In the process of PSO, each single solution in the D-dimensional search space is taken as a “bird” called “particle”. The *i*th particle position is represented as vector ***X***_*i*_ = (*x*_*i*1_,*x*_*i*2_,…,*x*_*iD*_)^*T*^. A particle is characterized by position, velocity and fitness value. The position giving the best fitness value of the *i* th particle is represented as vector ***P***_*i*_ = (*p*_*i*1_,*p*_*i*2_,…,*p*_*iD*_)^*T*^and the velocity is represented as vector ***V***_*i*_ = (*v*_*i*1_,*v*_*i*2_,…,*v*_*iD*_)^*T*^. The fitness value is calculated by fitness function and can display the pros and cons of a particle. The index of the best particle among all the particles in the population is represented as g. The particles are operated in accordance with the following equations: 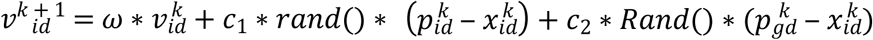 and 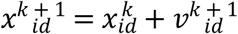. In the formulas, the *ω* is inertia weight, the k is the current iteration number, the *c*_1_and *c*_2_ are acceleration factor, the range of *i* is positive integer from 1 to n, the range of *d* is positive integer from 1 to D, the *Rand*() is random number value distributing between 1 and 2 [22, 30].PSO is initialized with a group of random particles (solutions) and then searches for optima by updating generations. The parameters of PSO were a population size of 20, and 200 iterations [24, 30]. The data in the Table 4 were an average of triplicate observations and subjected to one-way analysis of variance. The extraction solvents (petroleum ether and acetone) were set as 1 and 2, respectively. All calculations were implemented with Matlab 7.8.0.347 R2009a software (The MathWorks, Natick, MA, USA).

## Results and discussion

### The modeling of FAs

Crude fat content of the three categories of *Coix* seed and the statistical results of the calibration and validation set for the FA content are summarized in Table 1. The BCS had the highest crude fat content, which corresponded to the maximum average value of FAs content. The TCS showed the minimum crude fat content and resulted in the minimum of FAs content. There were significant differences in the FA content of the different categories of *Coix* seeds. The FA content of SCS had the maximum deviation in different extraction conditions, the BCS ranked second, and the TCS had the minimum. The maximum FA content of SCS (163.30 mg KOH/100 g) was 1.04 times that of BCS and 2.02 times that of TCS, while the minimum of BCS was 1.19 times that of SCS. It might be ascribed that the fat of SCS was tightly bound with starch, protein, phosphorus [31-33] and other nutritional ingredients, and inappropriate extraction conditions made it difficult to extract the fat. The FA concentration range of the validation set was located in the range of calibration set, which was suitable for acquiring successful calibrations.

**Table 1.** Statistical data for crude fat and fatty acids (FAs) of the three categories of *Coix* seed in the calibration and validation sets. Means followed by a different letter within a column for each *Coix* seed are significantly different (P < 0.05). BCS, big *Coix* seed; SCS, small *Coix* seed; TCS, translucent *Coix* seed; SD, standard deviation.

| Varieties | Crude fat (%) | Calibration set of FAs (mg KOH/100 g) |  |  |  | Validation set of FAs (mg KOH/100 g) |  |  |  |
| --- | --- | --- | --- | --- | --- | --- | --- | --- | --- |
|  |  | No. of samples | SD | Mean | Range | No. of samples | SD | Mean | Range |
| BCS | 7.68±0.04a | 24 | 12.41 | 140.07 | 116.10–157.60 | 8 | 13.50 | 143.44 | 121.40–157.40 |
| SCS | 7.10±0.08b | 24 | 23.00 | 132.55 | 97.20–163.30 | 8 | 25.25 | 125.07 | 99.50–160.50 |
| TCS | 5.45±0.15c | 24 | 11.39 | 65.42 | 42.70–80.80 | 8 | 12.41 | 66.54 | 51.30–80.10 |

The results of the PLSR and BPNN models for both the calibration and validation set of FAs is shown in Table 2. Generally, a good model should have a low RMSEP value and a high R^2^ value. Both modelling methods showed good performance for the three categories of *Coix* seed. However, by making a comparison between the PLSR and BPNN models, it can be seen that the PLSR model performances of BCS and SCS were slightly superior to those of the BPNN model which had a higher R^2^ (0.9299 and 0.9744, respectively), showing that the relationship between FA content and extraction conditions for BCS and SCS was more likely to be a linear function. While the BPNN model of TCS was a little better than that of the PLSR model, the R^2^ and RMSEP values were 0.8575 and 0.2981, respectively, which might indicate that the relationship of FA content and extraction conditions was more aligned with a nonlinear function. The results demonstrated that the three types of *Coix* seed had different extraction mechanisms. Although the SCS had the best model performance, the results for TCS still need to be improved. It is necessary to carry out more trials by adjusting the interval of extraction conditions (temperature and time) to improve the robustness and predictive ability of the PLSR and BPNN models in the future studies. And yet, the model performances of BCS, SCS, and TCS have proven that the PLSR and BPNN can reflect on the relationship of FA content and extraction conditions well and also rapidly and efficiently predict the FA content of *Coix* seed.

**Table 2.**
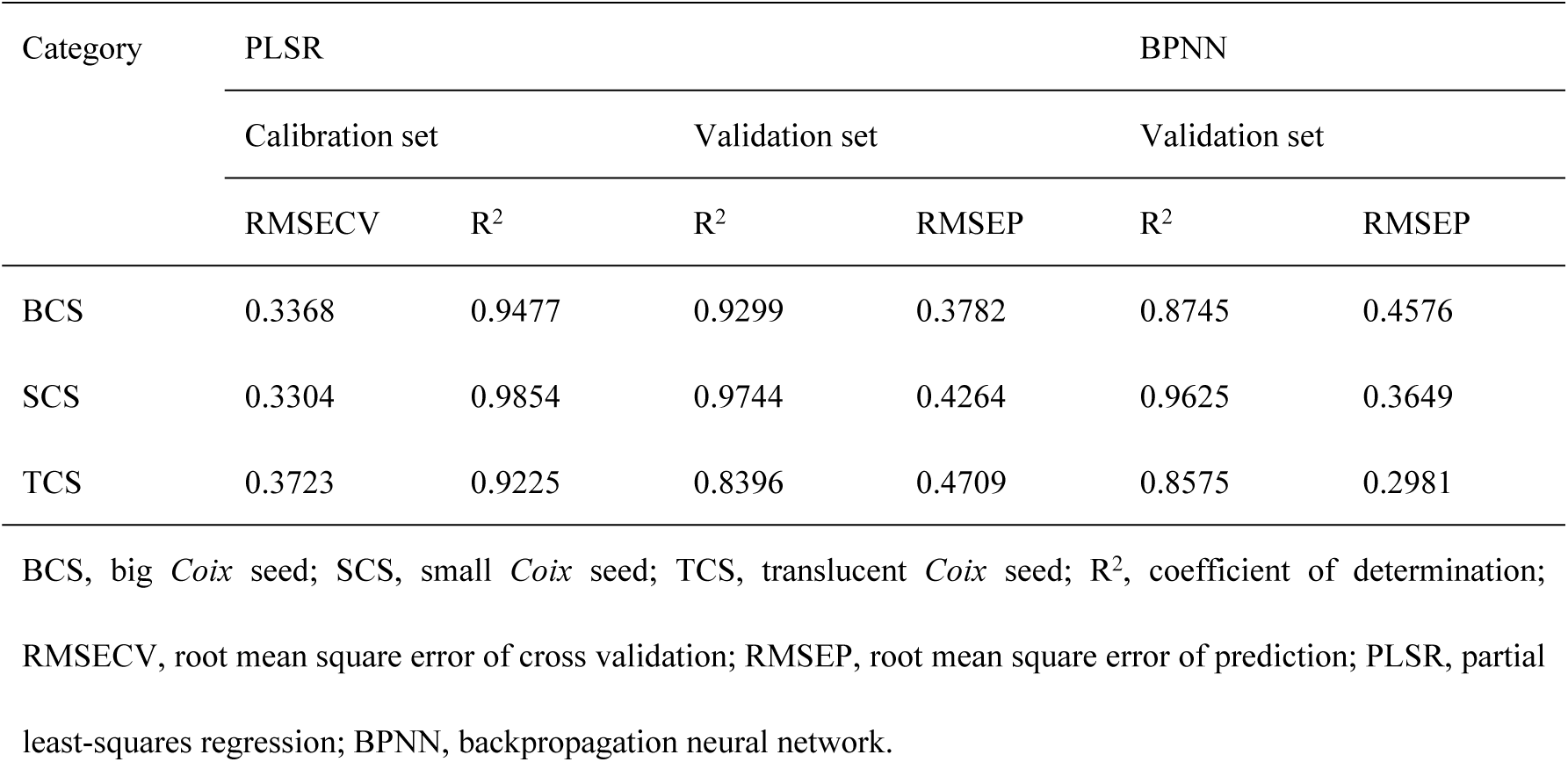
The performances of PLSR and BPNN models for the fatty acid content of the three categories of *Coix* seed. BCS, big *Coix* seed; SCS, small *Coix* seed; TCS, translucent *Coix* seed; R2, coefficient of determination; RMSECV, root mean square error of cross validation; RMSEP, root mean square error of prediction; PLSR, partial least-squares regression; BPNN, backpropagation neural network.

### Optimization of the FA extraction conditions

The fitting data of the optimal PLSR and BPNN models were used as a fitness function for GAs and PSO. The GAs and PSO searched for the optimal combination in the range of extraction conditions. The evolution process and optimization results of GAs and PSO are shown in Fig. 2 and Table 3. It can be seen from Fig. 2 that the PSO algorithm had faster convergence than that of GAs and the second iteration had obtained the optimum fitness values for the three types of *Coix* seed [22]. However, there were the problems of premature convergence, low precision, and low iterative efficiency in the PSO algorithm, which could result in a local optimum when tackling complex problems. Table 3 showed that the PSO algorithm could obtain the BCS and SCS trapped in the local optimum, because there was great difference between the actual and predicted FAs values for the same extraction conditions (130 °C, 20 min, and acetone extraction). For BCS and SCS, after 86 and 63 evolutional generations by GAs, respectively, the maximum theoretical FAs contents were 158.39 and 165.62 mg KOH/100 g, respectively. The optimal extraction conditions (rounded data) of BCS were 123 °C, 18 min, and acetone extraction (Because the predicted solvent values was 1.94, the rounded data was 2, which represented acetone solvent), and those of SCS were 126 °C, 20 min, and acetone extraction. For TCS, the optimal algorithm was PSO, and the predicted FAs content was 81.54 mg KOH/100 g with the extraction conditions (rounded data) of 124 °C, 20 min and acetone solvent. Then the FAs contents were determined again at the optimal extraction conditions obtained from the above chemometrics methods, and the results are displayed in Table 4. It could be seen from Table 4 that the actual FAs contents of the three *Coix* seeds (BCS was 160.12mg KOH/100g, SCS was 166.01mg KOH/100g and TCS was 81.28mg KOH/100g) in the optimal extraction conditions were approximated to the predicted values (BCS was 158.39mg KOH/100g, SCS was 165.62mg KOH/100g and TCS was 81.54mg KOH/100g), and higher than the highest FAs contents (BCS was 157.60mg KOH/100g, SCS was 163.30mg KOH/100g and TCS was 80.80mg KOH/100g) used in modelling. Furthermore, the results have also proved that the optimal extraction conditions are reasonable. The above results indicated that the extraction time of ASE (not higher than 1h) was significantly lower than that of supercritical fluid extraction (not lower than 2.5h) [8, 11]. The extraction efficiency of acetone for FAs in *Coix* seed was higher than petroleum ether, and the FAs in the BCS were the most easily extracted among the three types of *Coix* seed. The GAs and PSO were rapid and effective extreme value searching algorithms, although the classical PSO needs to be further improved.

**Table 3.** Comparison of the optimization extraction parameters by PLSR and BPNN combined with GAs and PSO. BCS, big *Coix* seed; SCS, small *Coix* seed; TCS, translucent *Coix* seed; PLSR, partial least-squares regression; BPNN, backpropagation neural network; GA, genetic algorithms; PSO, particle swarm optimization.

| Varieties | Optimization methods | Temperature (°C) | Time (min) | Solvent | FA (mg KOH/100 g) |
| --- | --- | --- | --- | --- | --- |
| BCS | PLSR-GA | 123.35 | 18.19 | 1.94 | 158.39 |
|  | PLSR-PSO | 130 | 20 | 2 (acetone) | 164.29 |
| SCS | PLSR-GA | 126.14 | 19.68 | 1.99 | 165.62 |
|  | PLSR-PSO | 130 | 20 | 2 (acetone) | 168.33 |
| TCS | BPNN-GA | 123.95 | 19.75 | 1.99 | 81.10 |
|  | BPNN-PSO | 124.38 | 20 | 2 (acetone) | 81.54 |

**Fig 2.**
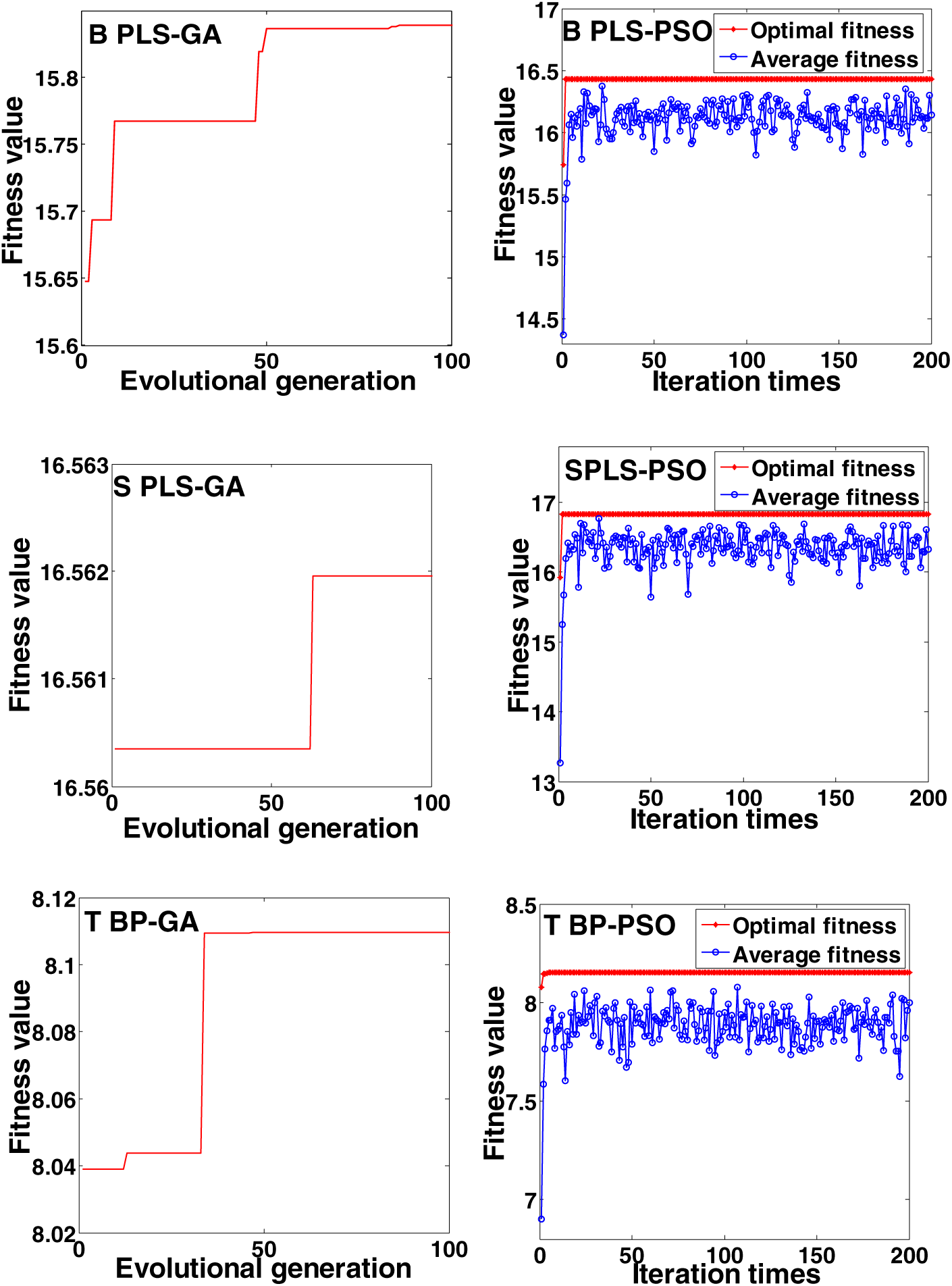
Evolution of the optimal and average fitness values in genetic algorithms and particle swarm optimization. The B, S, and T represent big *Coix* seed, small *Coix* seed, and translucent *Coix* seed, respectively.

### FA comparison of the three types of *Coix* seed oil

The FA composition and content of the three categories of *Coix* seed oil are displayed in Table 5. It is seen from Table 5 that the types of FAs were slightly different from those of supercritical extractions [8]. There were not heptadecenoic acid, nonadecanoic acid, nonadecyenoic acid, eicosenoic acid, heneicosanic acid, tricosanoic acid, tetracosanoic acid, and pentacosanoic acid in the study of Hu *et al*. [8],while eicosanoid and hexacosanoic acid were not detected in our study. Oleic acid accounted for the highest proportion in the oils from the three categories of *Coix* seed; the contents were BCS 75.26%, TCS 77.02%, and SCS 73.45%, respectively, which was higher than that of the previous study (47.5%) which used supercritical extraction [8]. These results could be due to the differences of extraction methods. From the Table 5, it could be seen that there were significant differences in the content of palmitic acid, palmitoleic acid, stearic acid, oleic acid, eicosanoic acid, and tetracosanoic acid among the three oils. Furthermore, the BCS showed a significant difference with TCS and SCS in the content of heptadecanoic acid, heptadecenoic acid, nonadecyenoic acid, eicosenoic acid, and docosanoic acid. That is, the FA composition of SCS was a little closer to that of TCS. The results explained that the FA composition of different varieties of *Coix* seed were different and could be ascribed to the differences in biological origin.

**Table 4.** Verification the FA content of three *Coix* seeds in the optimization extraction parameters. BCS, big *Coix* seed; SCS, small *Coix* seed; TCS, translucent *Coix* seed; PLSR, partial least-squares regression; BPNN, backpropagation neural network; GA, genetic algorithms; PSO, particle swarm optimization.

| Varieties | Optimization methods | Temperature (°C) | Time (min) | Solvent | FA (mg KOH/100 g) |
| --- | --- | --- | --- | --- | --- |
| BCS | PLSR-GA | 123 | 18 | acetone | 160.12±1.01 |
| SCS | PLSR-GA | 126 | 20 | acetone | 166.04±0.71 |
| TCS | BPNN-PSO | 124 | 20 | acetone | 81.28±0.95 |

**Table 5.** Fatty acid analysis of the three categories of *Coix* seed oil. Means followed by a different letter within a row for each *Coix* seed oil are significantly different (P < 0.05). BCS, big *Coix* seed; SCS, small *Coix* seed; TCS, translucent *Coix* seed.

| Fatty acid (%) | BCS | SCS | TCS |
| --- | --- | --- | --- |
| Myristic acid | 0.08±0.00a | 0.07±0.00a | 0.05±0.00b |
| Pentadecyl acid | 0.04±0.00a | 0.04±0.00a | 0.03±0.00a |
| Palmitic acid | 16.25±0.07b | 16.80±0.04a | 15.04±0.06c |
| Palmitoleic acid | 0.65±0.01b | 0.75±0.01a | 0.51±0.01c |
| Heptadecanoic acid | 0.18±0.01b | 0.25±0.01a | 0.25±0.01a |
| Heptadecenoic acid | 0.14±0.01b | 0.18±0.00a | 0.21±0.01a |
| Stearic acid | 4.19±0.02b | 5.35±0.01a | 3.96±0.03c |
| Oleic acid | 75.26±0.06b | 73.45±0.04c | 77.02±0.03a |
| Octadecadienoic acid | 0.09±0.00a | 0.10±0.00a | 0.09±0.00a |
| Nonadecanoic acid | 0.03±0.00a | 0.04±0.00a | 0.03±0.00a |
| Nonadecyenoic acid | 0.06±0.01b | 0.07±0.00a | 0.09±0.01a |
| Eicosanoic acid | 1.28±0.01b | 1.31±0.01a | 1.18±0.01c |
| Eicosenoic acid | 0.81±0.01b | 0.84±0.01a | 0.86±0.01a |
| Heneicosanoic acid | 0.03±0.00a | 0.03±0.00a | 0.03±0.00a |
| Docosanoic acid | 0.44±0.00a | 0.34±0.01b | 0.34±0.00b |
| Docosenoic acid | 0.04±0.00a | 0.02±0.00a | 0.04±0.00a |
| Tricosanoic acid | 0.05±0.00a | 0.05±0.00a | 0.04±0.00a |
| Tetracosanoic acid | 0.38±0.00a | 0.29±0.01b | 0.21±0.01c |
| Pentacosanoic acid | 0.02±0.00a | 0.02±0.00a | 0.01±0.00a |

## Conclusions

The results demonstrated that the PLSR and BPNN models could reflect the relationship of FA content and extraction conditions well. For BCS and SCS, the performances of the PLSR models slightly outperformed those of the BPNN models; while for TCS, the BPNN model was superior to the PLSR model. The GAs could seek out the optimal extraction conditions for the PLSR models of BCS (123 °C, 18 min, and acetone extraction) and SCS (126 °C, 20 min, and acetone extraction) rapidly and effectively, and PSO algorithms were more suitable for the BPNN model of TCS (124 °C, 20 min, and acetone extraction). Furthermore, all the extraction time of the FAs from three *Coix* seeds using ASE was shorter than common extraction techniques, such as Soxhlet extraction, microwave extraction, sonication extraction and supercritical fluid extraction. There were differences in the FA content of the three categories of *Coix* seed on account of the differences of biological origin. Therefore, ASE combined with chemometrics methods can be a labour-saving, time-saving, and powerful tool for rapid and effective determination of FAs compared with the common extraction methods. We believe that this approach should be further applied to extract other nutrition ingredients from natural food samples.

## Acknowledgements

This work was supported by National Natural Science Foundation of China (31772189 and 31171642) and The Youth Talent Development Plan of Shanghai Agriculture Committee of China [Grant no. 2017(1-31)].

